# Neurophysiological correlates of passive movements are speed- and type-dependent

**DOI:** 10.1101/2025.07.09.663910

**Authors:** M.P. Veldman, J.Z. Kwant, J. Lommerse, M. Feenstra, C.J.C. Lamoth, A.H.M. Volkers, H. Drenth, S. Zuidema, I. Bautmans, H. Hobbelen

## Abstract

**Introduction:** The supraspinal involvement in the control of passive movements remains elusive. Mechanoreceptor properties, their change in the context of ageing and the somatotopically organized supraspinal connections between sensory and motor systems provide a neuroanatomical basis for the prediction that cortical structures are involved in the control of passive movements. Previous electromyographic evidence indeed show movement speed and - type-dependent changes in muscle activity. This study aimed to provide electrophysiological evidence for the involvement of frontal cortex inhibition and corticomotor interactions in the control of passive movements.

**Methods:** Continuous and discontinuous passive elbow movements were performed in healthy younger (n = 20, 22.5 ± 2.31 y) and older (n = 20, 72.7 ± 5.73 y) adults at three movement speeds (20, 60, and 100 bpm) while electro-encephalographic (EEG) and electromyographic (EMG) data were acquired. Alpha power and beta corticomuscular connectivity were used as measures of frontal cortex inhibition and brain-muscle connectivity, respectively.

**Results:** Frontal cortex inhibition decreased (p = 0.036) and brain-muscle connectivity increased (p < 0.001) with increasing movement speeds. In addition, frontal cortex inhibition was 17% higher in the discontinuous condition as compared to the continuous condition (p = 0.005) while corticomuscular coherence was 25.9% higher in the continuous vs. the discontinuous condition (p < 0.001). These effects were independent of age.

**Conclusion:** The present results provide insights into the control of passive movements and show that frontal cortex inhibition and brain-muscle interactions depend on movement speed and movement type.

## 1. Introduction

Passive motion influences human functioning at various levels. At the muscle architecture level, muscle fibers are combined to whole muscles by connective tissue that passively moves along with contractions. The structural organization of connective tissue therefore substantially influences in the magnitude of force that muscles exert during contractions (Enoka, 2015). Changes in muscle length during contractions – induced by both active and passive movements – are detected by various mechanoreceptors, that in turn affect bodily functions. Stretch-sensitive type III mechanoreceptors play a role in increasing ventilatory and cardiovascular responses at the organ level [for a review, see (Trinity and Richardson, 2019)] as well at as the muscle level. Within muscles, muscle spindles detect (the rate of) change in muscle fiber length and send afferent volleys to the spinal cord via type Ia and II afferent nerves that may trigger spinal reflexes (Enoka, 2015). These sensory afferents ascend to the primary sensory cortex and form direct, somatotopically organized connections with the primary motor cortex [for a review, see (Veldman et al., 2014)]. This provides a neuroanatomical basis for influence of passive movements on supraspinal processes.

Passive movements indeed elicit cortical activity. For example, conjunction analyses of functional imaging data revealed overlap in activity in sensorimotor and cerebellar areas when healthy young individuals performed active and passive pedaling movements [e.g., (Jaeger et al., 2014)]. Since mechanoreceptors detect rates of change, and their inputs reach supraspinal structures, this provides neuroanatomical and neurophysiological support that cortical activity varies with different passive movement speeds. Moreover, since muscle spindle sensitivity decreases with advancing age (Lord et al., 2018), the effect of passive movements on cortical activity may also change with age. Indirect support for this hypothesis comes from a series of studies by Marinelli and colleagues, suggesting an age- and movement-speed-dependent association between passive movements and cortical activity. They observed involuntary increases in electromyographic (EMG) activity in healthy individuals, with these increases being more pronounced in older age groups and during fast, continuous movements (Marinelli et al., 2022, 2017). Notably, this phenomenon further increased in individuals with cognitive decline, offering insights into the mechanisms underlying paratonia, a dementia-related motor symptom that is characterized as an involuntary resistance to passive movements. A prominent hypothesis postulates that pathological responses to passive movements may originate from frontal cortex dysfunction (Beversdorf and Heilman, 1998; Drenth et al., 2020), which is in line with data showing that motor output is under inhibitory control of the prefrontal cortex (Aron et al., 2004) and that passive movements elicit activity in frontal areas (Jaeger et al., 2014).

Direct neurophysiological evidence for this hypothesis, however, is missing. In this study, we use electroencephalography (EEG), that measures oscillatory brain activity at the scalp with a high temporal resolution, to quantify frontal cortex inhibition and brain-muscle connectivity. Specifically, previous research showed that oscillations in the alpha frequency range (8 – 12 Hz) reflect inhibitory processes (Klimesch et al., 2007). Moreover, because oscillations at the beta frequency range have been shown to be implicated in motor processes (Pfurtscheller and Lopes da Silva, 1999; van Wijk et al., 2012), the coherence between EEG signals from electrodes located over the motor cortex and EMG signals, i.e., corticomuscular connectivity, reflects brain-muscle interactions and is sensitive to age-related changes [e.g., (Spedden et al., 2019)].

The present study aims to provide direct electrophysiological evidence for the hypothesis that neurophysiological responses to passive movements are age- and speed-dependent. Based on properties of mechanoreceptors and previous findings in the context of paratonia (Marinelli et al., 2022, 2017), we hypothesized that (1) frontal cortex inhibition decreases with increasing age and movement speed and is lower in continuous vs. discontinuous movements based on elevated EMG-levels during such movements in a previous study (Marinelli et al., 2017). Because other data show that higher muscle tone in people with spasticity occurs in absence of increases in fusimotor drive from the spinal cord suggestive of disinhibition [for a reviews, see (Nordin et al., 2017; Sheean, 2002)], we also hypothesized that (2) brain-muscle connectivity increases with age and movement speed, and that the magnitude of such connectivity is higher in continuous vs. discontinuous movements.

## 2. Methods

### 2.1 Participants

Healthy young (n = 20, 22.5 ± 2.31 y) and older (n = 20, 72.7 ± 5.73 y) adults participated in this study. The sample size was based on a priori power analysis for repeated measures ANOVA (effect size: 0.25; power: 0.8). Participants were recruited from Groningen and the surrounding area by local flyers and mouth-to-mouth advertising. They were included if they were free of neurological disorders or other physical and psychological disorders and did not take drugs that affect nerve conduction velocity or cognitive processes, such as antidepressants, antipsychotics or sleep medications. Participants provided written informed consent before participating in a protocol that was approved by the Medical Ethical Review Committee (METc, NL81562.042.22) of the University Medical Center Groningen (UMCG) and was conducted according to the Declaration of Helsinki (2013).

### 2.2 Experimental design

This study adopted a cross-sectional design based on previous work (Marinelli et al., 2017) that was extended with synchronous EEG and EMG recordings. Baseline measures included anthropometrics, handedness [Edinburgh Handedness Inventory, EHI (Oldfield, 1971)], physical [Timed Up and GO, TUG (Podsiadlo and Richardson, 1991)] and cognitive functioning [Montreal Cognitive Assessment, MoCA (Nasreddine et al., 2005)]. In addition, the Paratonia Assessment Instrument (PAI) was performed to subjectively assess the presence of paratonia by interpreting the extent of resistance during passive movements (Hobbelen et al., 2008).

### 2.3 Experimental protocol

The experimental protocol started with a seated control task after which a series of arm movements with the right arm followed while brain (EEG) and muscle (EMG) activity was recorded. During the experiment, participants sat on a chair with their knees flexed at a ninety-degree angle and their feet flat on the ground. Participants performed passive arm movements with the dominant right arm at three different speeds, i.e., 20, 60 and 100 beats per minute (Figure 1). The participants were instructed to relax their arm during the passive arm movements while the researcher moved the arm through its full range thirty times (i.e., thirty flexion and thirty extension movements). At all three speeds, the passive movement conditions were performed in a continuous (i.e., one beat of the metronome between maximal flexion and maximal extension) and discontinuous (i.e., one beat of the metronome between maximal flexion and maximal extension and one beat of the metronome of pause between the end of a flexion/extension movement and the start of the next extension/flexion movement) fashion, resulting in a total of six conditions. All arm movements were performed on the rhythm of a metronome (Tempo Lite, Frozen Ape Pte. Ltd.), which was exclusively audible to the researcher through earphones. The order of the tasks was pseudorandomized so that half of the participants started with the continuous movements and the other half with the discontinuous movements.

**Figure 1.**
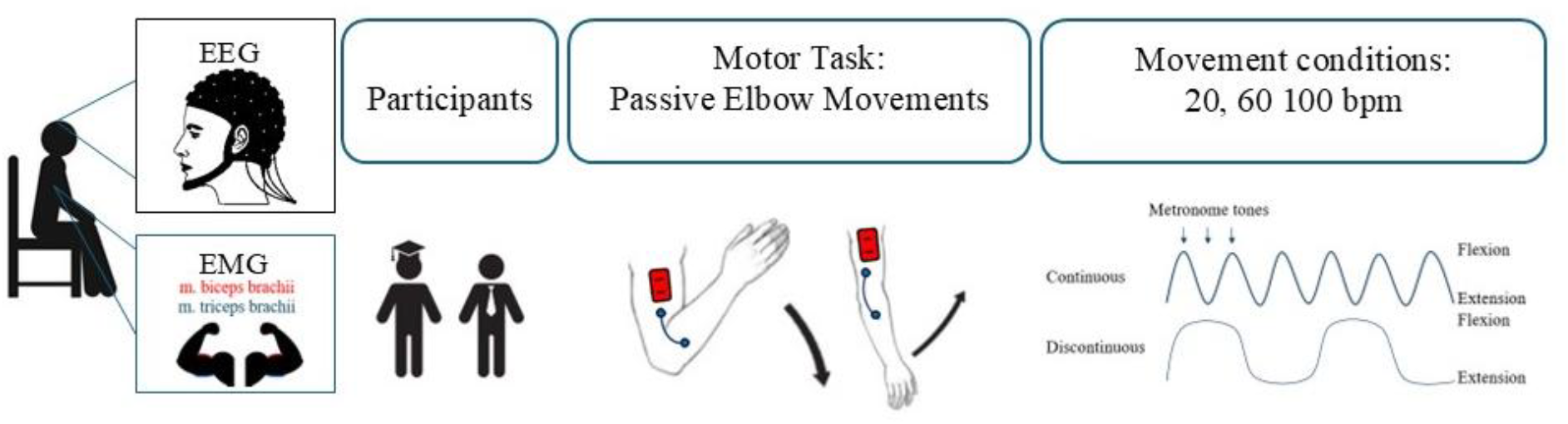
Overview of the experimental design. In healthy younger (n = 20) and older (n = 20) individuals, continuous and discontinuous passive movements were performed at three different speeds while 64-channelelectroencephalography (EEG)and biceps and triceps electromyography (EMG) data were acquired. bpm, beats per minute.

**Figure 1.**
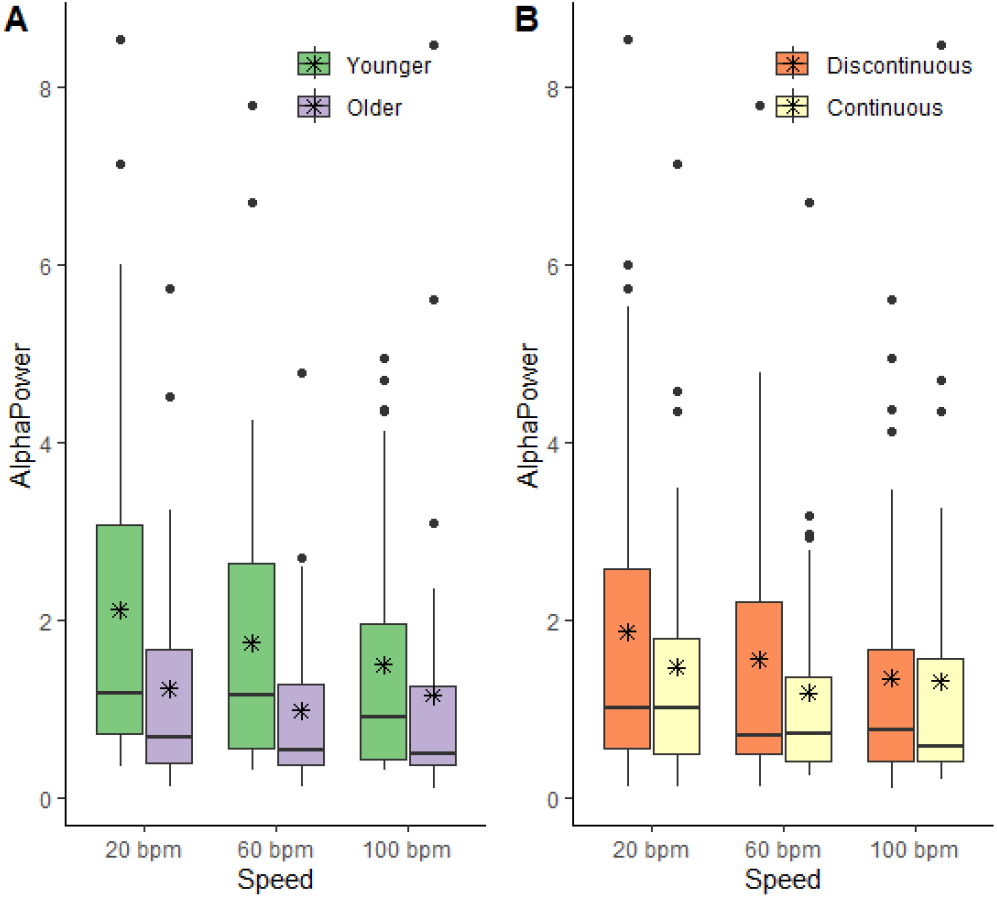
Effects of age, movement speed and movement type on alpha power. The panels display the effects of speed (panels A-B) and movement type (panel B) on alpha power. The horizontal lines and asterisks within the boxes represent the median and mean for each group/condition, respectively. Bpm, beats per minute.

Breaks were allowed depending on the participant’s need for rest. Figure 1 provides a schematic overview of the experimental design.

### 2.4 Electrophysiological data acquisition and analysis

#### 2.4.1 Data acquisition

Muscle activity from the right M. biceps brachii and M. triceps brachii were recorded using surface wireless EMG sensors (Trigno Avanti, Delsys, Natick, MA, USA; 27 x 37 x 13 mm). The EMG sensors were placed over the right biceps and triceps brachii muscles, which were determined by palpating the muscle during voluntary muscle contraction. EMG data were recorded at a sampling frequency of 1926 Hz.

Electroencephalography (EEG) data were continuously acquired with a 64-channel EEG system with Ag-AgCl electrodes placed on the scalp according to the international 10-20 system (TMSi SAGA64+, Oldenzaal, the Netherlands). The EEG data were acquired at 2048 Hz with an average reference. The impedance was kept below 10 kΩ. To minimize noise in the EEG signals, participants were asked to not speak, to avoid head movements, to relax the jaw and neck muscles and to minimize eye blinking during the measurements. A hard-wired trigger (0-5 V ramp-up TTL trigger) ensured the synchronous acquisition of EEG and EMG data.

#### 2.4.2 Data preprocessing

The EEG data were pre-processed in MATLAB R2022b (MathWorks), using scripts based on FieldTrip (Oostenveld et al., 2011). The data were filtered with a 3 Hz high-pass filter (6^th^ order Butterworth) and a 70 Hz low-pass filter (6^th^ order Butterworth). To remove electromagnetic line noise, a notch filter with a frequency of 50 Hz and its second and third harmonic (100 and 150 Hz) was also applied. After resampling the data to 1024 Hz, EEG channels were visually inspected and empty channels or channels with substantial artifacts were removed. Next, an independent component analysis was performed to remove artifacts caused by eye movements and muscle contractions based on time series and topographical distributions of power. This process was repeated for each condition and consistently performed by the same researcher (AHMV). Three components per trial were removed on average. The preprocessing pipeline concluded with referencing the data to an average reference.

The EMG data was pre-processed in MATLAB R2022b (MathWorks). First, the EMG data were restructured in the FieldTrip format to enable the preprocessing of the EMG data using similar procedures as for the EEG data. The EMG data were bandpass filtered using a 4^rd^ order Butterworth filter between 4 and 100 Hz. Additionally, electromagnetic line noise was removed with a 4^rd^ order Butterworth notch filter of 50 Hz and its second and third harmonic (100 and 150 Hz). The EMG data were full-wave rectified using an absolute Hilbert transform and resampled to 1024 Hz.

#### 2.4.3 Data analysis

The preprocessed EEG time series were epoched into non-overlapping 1-s-long epochs before being transformed to the frequency domain using a multitaper Fast Fourier Transformation with a Hanning window leading to a 1 Hz frequence resolution. Power in the alpha frequency range (8 – 12 Hz) was averaged for five frontal electrodes [F3, Fz, F4, FC3 and FC4; (Gompf et al., 2017)].

To quantify brain-muscle interactions, EEG and EMG data were concatenated and epoched in non-overlapping 1-s-long segments before being transformed to the frequency domain between 1 and 40 Hz using a multitaper Fast Fourier Transformation method with 5 Hz smoothing. Subsequently, corticomuscular coherence was calculated using the auto- and cross spectra (Liu et al., 2019). Then, beta-range (13 – 30 Hz) corticomuscular coherence was estimated between the pre-processed EEG timeseries in the electrode positioned over the upper arm region of the primary motor cortex contralateral to the right arm, represented by the C3, and the pre-processed EMG timeseries recorded from the biceps and triceps muscles. For each condition, the level of corticomuscular coherence was quantified as the area under the curve between the corticomuscular coherence estimates and significance line of the amplitude spectrum in the frequency domain (Halliday et al., 1995).

### 2.5 Statistical analysis

Statistical analyses were performed in SPSS (Version 26, IBM, Chicago, IL, USA). The Shapiro-Wilk test revealed that all data were normally distributed. Descriptive statistics (in means and standard deviations) were presented for baseline age, physical and cognitive performance, and handedness by age group. To examine whether age, movement speed and movement type impacted alpha power data and beta corticomuscular coherence, data were analysed with repeated measurement multivariate analysis of variance (rmMANOVA) for each hypothesis. In these rmMANOVA’s, movement speed (20, 60 and 100 bpm) and type of movement (continuous and discontinuous) were included as within-subject factors and age (healthy younger and older) was included as between-subjects factor. The significance level of the Box’s M test was set at p <.001, as this test is considered highly sensitive (Jiamwattanapong and Ingadapa, 2021). When the assumption of sphericity was violated as evidenced by the Mauchly’s test, a Greenhouse-Geisser correction of degrees of freedom was used. Partial eta square 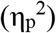 was used as a measure of effect size where ≥ 0.2, ≥ 0.5 and ≥ 0.8 were considered as small, medium and large effects, respectively (Cohen, 2013). For all analyses the level of significance was set at p < 0.05.

## 3. Results

Seventeen older and twenty younger individuals completed this study. Three of the included older adults were excluded from further participation because they scored <26 on the MoCA. Subjective paratonia was observed in two older adults and not in younger adults. Table 1 provides an overview of the baseline measures.

**Table 1.** Descriptive statistics

|  | Older adults (n = 20) | Younger adults (n = 20) |
| --- | --- | --- |
| Age in years (mean ± SD) | 72.7 ± 5.73 | 22.5 ± 2.31 |
| MoCA<sup>1</sup> (mean ± SD) | 26.1 ± 2.6 | 27.6 ± 1.7 |
| TUG<sup>2</sup> (s) in seconds (mean ± SD) | 7.87 ± 1.09 | 6.8 ± 1.44 |
| EHI<sup>3</sup> | 96.7 ± 5.9 | 66.2 ± 54.3 |

Abbreviations: MoCa, Montreal Cognitive Assessment, TUG, Timed Up and Go; EHI, Edinburgh Handedness Inventory.
- 1. Higher scores indicate better performance - 2. Higer scores indicate worse performance - 3. Higher scores indicate stronger right-handedness

Alpha power decreased with increasing movement speeds (F_(1.558, 52.978)_ = 3.91, p = 0.036, 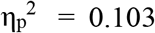 Figure 1A-B) and was 17% higher in discontinuous as compared to continuous movements (F_(1, 34)_ = 8.85, p = 0.005, 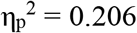 Figure 1B). There was no effect of age (F_(1, 34)_ = 1.262, p = 0.269, 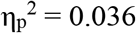) and none of the interaction effects reached significance.

Corticomuscular coherence between the primary motor cortex and the biceps muscle increased with increasing movement speed (F_(2, 26)_ = 108.04, p < 0.001, 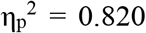 Figure 2A-B).

**Figure 2.**
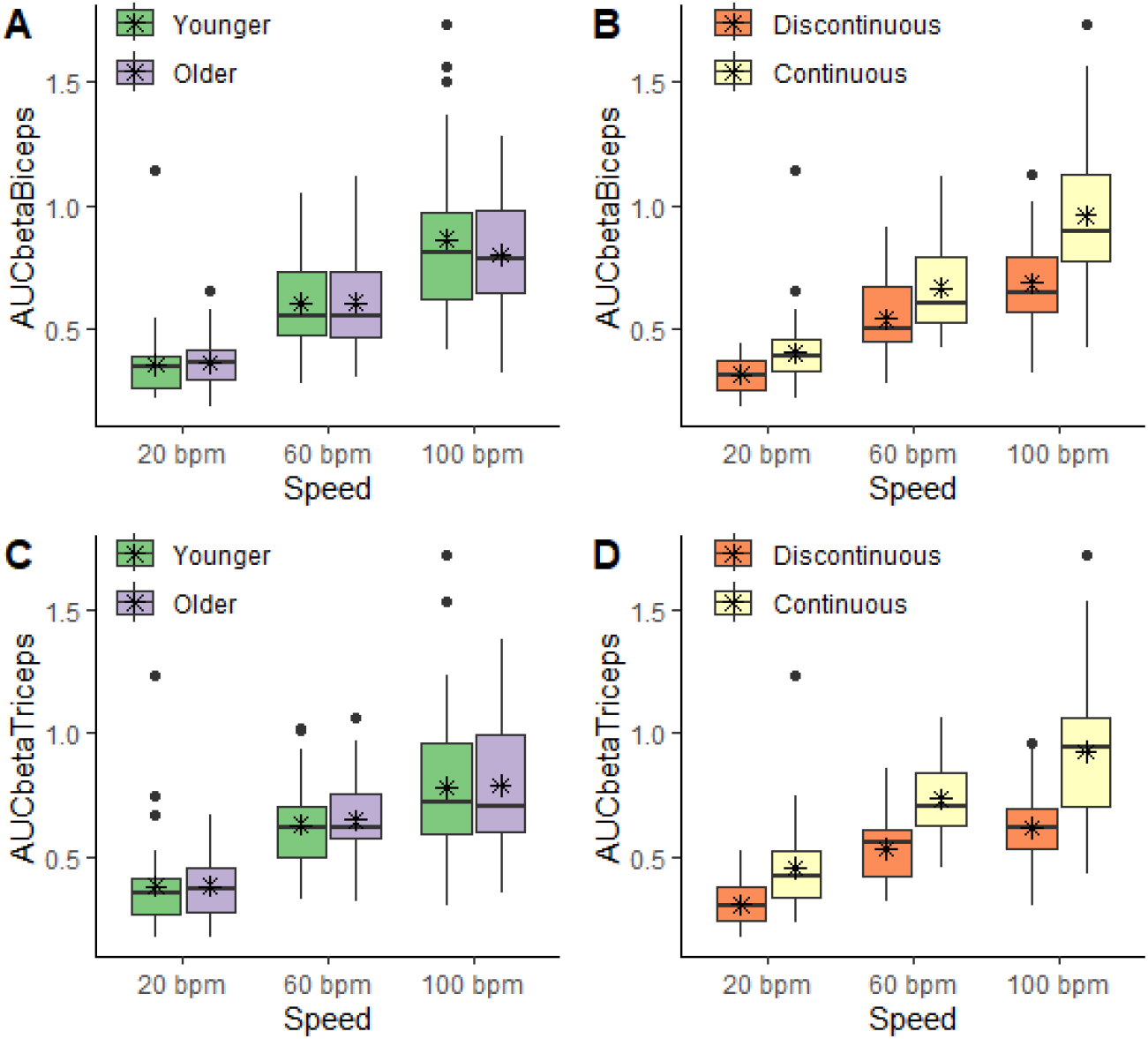
Effects of age, movement speed and movement type on corticomuscular coherence. The panels display the effects of speed (panels A-C) and movement type (panels B-D) on corticomuscular coherence between the primary motor cortex and the biceps muscle (panels A-B) and triceps muscle (panels C-D). The horizontal lines and asterisks within the boxes represent the median and mean for each group/condition, respectively. Bpm, beats per minute.

Additionally, the continuous condition showed 25.9% higher corticomuscular coherence as compared to the discontinuous condition (F_(1, 27)_ = 27.08, p < 0.001, 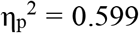 Figure 2B). Similarly, corticomuscular coherence between the primary motor cortex and the triceps muscle increased with increasing movement speed (F_(1.654, 44.663)_ = 77.26, p < 0.001, 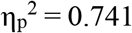 Figure 2C-D) and was 32.1% higher in continuous vs. discontinuous movements (F_(1, 27)_ = 40.38, p < 0.001, 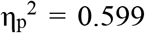 Figure 2D). There were no effects of age (Figure 2A-C) or interaction effects.

## 4. Discussion

This study examined the effects of age, movement speed and movement type on frontal cortex inhibition and brain-muscle connectivity during passive elbow flexion and extension movements. In line with the hypotheses, frontal cortex inhibition decreased and brain-muscle connectivity increased with increasing movement speeds. In addition, discontinuous movements were associated with higher inhibition and lower brain-muscle connectivity as compared to continuous movements. In contrast with our hypothesis, however, age did not affect frontal cortex inhibition and brain-muscle connectivity. The present study altogether reinforces the hypothesis of supraspinal involvement in the control of passive movements as indicated by movement speed and -type-dependent modulation of frontal cortex inhibition and brain-muscle connectivity.

### 4.1 Movement speed

The decrease in frontal cortex inhibition and increase in brain-muscle connectivity with increasing movement speeds are in line with our hypothesis and consistent with other studies showing decreased alpha power over sensorimotor regions with increasing speeds in a walking task (Nordin et al., 2020). These data suggest that the interplay between sensory inputs and motor outputs is important during the control of passive movements. It is likely that input from muscle spindles that detect the (rate of) change in muscle fiber length (Enoka, 2015), and reaches the primary and motor cortex via somatosensory afferents and somatotopic connections between the primary sensory and motor cortices (Veldman et al., 2014), played a role in the current findings. Imaging data suggest that the overlap in neural activation between voluntary and passive movements exceeds the primary sensorimotor regions and extends to frontal regions (Dobkin et al., 2004; Jaeger et al., 2014). Interestingly, peak activations during passive movements are generally weaker as compared to voluntary movements, except for activations in the secondary motor cortex (Dobkin et al., 2004), that has been suggested to be a mediating structure between sensory and motor areas based on animal data (Barthas and Kwan, 2017). Such interactions between sensory and motor systems may also explain the decreased inhibition in the frontal cortex. Specifically, decreased inhibition allows for increased responsiveness to incoming sensory information associated with faster movements (Klimesch, 2012; Mathewson et al., 2011), which may have triggered the increases in connectivity [(Jones, 1983; Terao et al., 1999), for a review see (Veldman et al., 2014)]. This suggestion is further supported by data showing event-related desynchronization in the beta range (Alegre et al., 2002), which is differently modulated depending of the movement phase, i.e., preparation, execution, termination [e.g., (Heinrichs-Graham et al., 2017; Pfurtscheller and Lopes da Silva, 1999)]. In summary, as passive arm movement speed increases, frontal cortex inhibition decreases and brain-muscle connectivity increases, possibly via a complex interplay between sensory afferents and motor efferents.

### 4.2 Continuous vs. discontinuous movements

The higher frontal cortex inhibition and reduced brain-muscle connectivity during discontinuous, as compared to continuous, movements may reflect the increased need to control motor output in discontinuous movements. The nature of the discontinuous movements requires repeated stopping and starting of an ongoing process. In this context, muscle spindles, which respond to the rate of change of muscle fiber length and subsequently may result in reflexive contraction of the muscle via the muscle spindle reflex arc (Macefield and Knellwolf, 2018), are arguably more active in discontinuous movements in comparison with continuous movements. Since participants were instructed to relax their muscles during the passive movements, we speculate that the increased alpha power in the frontal cortex could reflect active inhibition of this muscle spindle reflex arc that may have been necessary to prevent the reflexive contraction that is required for allowing passive movements.

The interpretation of more active inhibition during discontinuous movements would logically lead to the prediction that brain-muscle connectivity is lower in discontinuous as compared to continuous movements. The present study supports this prediction as the magnitude of brain-muscle connectivity was lower in discontinuous versus continuous movements. These findings are consistent with previous data indicating modulations of brain-muscle connectivity following passive movements in a walking neurorehabilitation context (Artoni et al., 2023), suggesting an overlap between neurophysiological underpinnings of active and passive movements. Moreover, because beta-range corticomuscular coherence has been implicated in the coordination of activity between different muscles (Reyes et al., 2017), the higher corticomuscular coherence in the continuous movement condition might reflect a higher degree of coordination between biceps and triceps activity in this condition. This interpretation is further supported by the absence of differences between the M. biceps and M. triceps brachii on brain-muscle connectivity. Altogether, the current and previous data indicate that supraspinal structures are more involved in continuous passive movements as compared to discontinuous movements.

### 4.3 Age

Contrary to the hypothesis, the modulation of frontal cortex inhibition and brain-muscle connectivity was independent of age. Previous studies revealed inconsistent age-related modulations in beta-range brain-muscle connectivity, with some showing age-related increases (Johnson and Shinohara, 2012; Kamp et al., 2013) while others reported age-related decreases in beta-range CMC (Bayram et al., 2015). Our hypothesis that such age-related differences would be present during passive movements was based on the deterioration of muscle spindle sensitivity with advancing age (Lord et al., 2018) and inspired by previous data showing agerelated increases in the presence of paratonia (Marinelli et al., 2022, 2017). Participants included in the final analyses, however, did not show signs of paratonia as assessed by the subjective paratonia assessment instrument (Hobbelen et al., 2008). We speculate that the interactions between sensory and motor systems are relatively robust against slight reductions in sensory function and that more advanced sensory deteriorations are required to detect agerelated changes at the cortical level. In this context, it is noteworthy that of the three participants who were excluded from the analyses because they scored <26 on the the MoCA, two showed slight to moderate signs of paratonia based on the subjective paratonia assessment instrument. If the current assumption that paratonia is mediated by motor cortex disinhibition as a result of frontal cortex dysfunction in people with dementia is true, the absence of age-related effects in the present sample is not surprising.

### 4.4 Clinical implications

The control of passive movements is affected in a number of neurodegenerative pathologies. In Parkinson’s disease, the unintentional display of heightened muscle tone in response to passive movements, hypertonia and “lead-pipe” rigidity, can only partly be attributed to short- and long-latency stretch reflexes (Kwon et al., 2014; Xia et al., 2011), which suggest a cortical origin of these symptoms. Indeed, in people with spasticity, heightened muscle tone is observed in the absence of increased fusimotor drive from the spinal cord suggestive of cortical disinhibition [for a review, see (Nordin et al., 2017)]. The current study design was based on previous studies investigating paratonia, the unintentional resistance to passive movements in people with dementia (Marinelli et al., 2022, 2017). Such impaired motor control has a severe impact on daily life, not only in performing activities of daily living but also in the care of individuals who can no longer actively manage these tasks themselves. Moreover, our data are consistent with electrophysiological data in people with paratonia that display more muscle activity at higher movement speeds and during continuous movements as compared to discontinuous movements. Furthermore, the current findings provide initial insights and support for the hypothesis that frontal cortex disinhibition play a role in the increased resistance to passive movements (Drenth et al., 2020). Future studies in people with dementia that show signs of paratonia can corroborate the hypothesis that frontal cortex dysfunction cause diminished control over passive movements. Together, these studies may pave the way for the development of targeted interventions for the treatment of pathological muscle tone.

### 4.5 Limitations

While the experimental design systematically evaluated the effect of passive movement speed and type and the passive movements were induced by trained experimenters, the personal approach may have resulted in slight variations in movement speed and amplitude. It is possible that these variations triggered involuntary movements or resistance to movements and consequently affected our neurophysiological outcome measures. Moreover, because we focused on the cortical and corticomotor correlates of passive movements, we opted to not control for the amplitude of the acquired EMG signals. Although alpha power is likely insensitive to the magnitude of the EMG signals and corticomuscular coherence is a normalized outcome measure, it would be interesting for future studies to evaluate whether alpha power in the frontal cortex and beta corticomuscular coherence are modulated with varying EMG levels. Lastly, as depicted in figures 2 and 3, the EEG-derived outcome measures, alpha power and beta brain-muscle interactions, displayed considerable variation, which may have reduced the statistical power and obscured age-related differences.

## 5. Conclusion

This study evaluated whether the neurophysiological correlates of passive movements depend on age, movement speed and type. The results revealed that, independent of age, frontal cortex inhibition decreased and brain-muscle connectivity increased with increasing movement speeds and that continuous, as compared to discontinuous, movements were associated with lower frontal cortex inhibition and higher brain-muscle connectivity. While future studies in patient populations are necessary to corroborate the prevalent hypothesis that frontal cortex function is involved in conditions that involve pathological muscle tones, the current data provide direct electrophysiological evidence that neurophysiological responses to passive movements are speed- and type-dependent.

## Acknowledgements

This study was supported by funds attributed to the International Joint Research Group ‘Move In-Age’ by the Vrije Universiteit Brussel (grant number: OZR3807). The authors wish to specifically acknowledge prof. dr. Sytse Zuidema who provided invaluable contributions to the development of the project and manuscript and sadly passed away on April 25th, 2025.

## Notes

### Competing Interest Statement

The authors have declared no competing interest.

